# Centromeric α-satellite DNA is a hotspot of genotoxic damage, incomplete repair, and cytoplasmic mislocalization

**DOI:** 10.64898/2026.05.01.722289

**Authors:** Azait Imtiaz, Mohammad Waseem, Hudson O’Neill, Christine M. Wright, Bo-Ruei Chen, Wioletta Czaja, Rafael Contreras-Galindo

## Abstract

Centromeric α-satellite DNA constitutes a highly repetitive and structurally specialized component of the human genome, yet the mechanisms underlying its damage susceptibility and repair fidelity under genotoxic stress remain undefined. Here, we demonstrate that genotoxic stress preferentially targets active centromeres, generating DNA double-strand breaks (DSBs) within α-satellite arrays. Using bleomycin as a defined genotoxic perturbation, we identify dynamic alterations in centromeric repeat content, manifesting as net copy number losses and gains across multiple chromosome-specific α-satellite arrays following damage. Similar centromere-associated damage signatures are observed in fibroblasts from patients with limited cutaneous systemic sclerosis, indicating that these features extend beyond experimental systems. Centromeric DSBs engage ATM-dependent DNA damage signaling and are repaired predominantly through RAD51-associated homologous recombination; however, repair fails to fully restore centromeric integrity. This incomplete repair is associated with defects in kinetochore organization, chromosome missegregation, and the formation of micronuclei containing centromeric DNA. Notably, ∼30% of these structures retain CENP-B but lacks detectable CENP-A, indicating disruption of centromere chromatin organization. Centromeric chromatin is frequently mislocalized to the cytoplasm following nuclear envelope perturbation, where immunofluorescence analysis reveals proximity to MHC class II (HLA-DRB1). Together, these findings establish centromeric α-satellite DNA as a vulnerability hotspot under genotoxic stress, with implications for chromosome instability and chromatin antigen exposure in fibrosis-associated autoimmunity.

## Introduction

Centromeres are essential genomic elements that ensure accurate chromosome segregation, yet their highly repetitive α-satellite DNA and specialized chromatin organization present unique challenges for DNA replication and repair (Aldrup-Macdonald and Sullivan 2014; McNulty and Sullivan 2018; Black and Giunta 2018). Human centromeres are composed of ∼171-bp α-satellite monomers arranged into higher-order repeat (HOR) arrays, which vary extensively in size and organization across individuals (Aldrup-Macdonald and Sullivan 2014; Logsdon et al. 2025). Despite recent advances in long-read genome assemblies, the susceptibility of centromeric repeats to genotoxic stress and the mechanisms governing their repair are incompletely characterized (Logsdon et al. 2025; Nassar et al. 2023). Emerging evidence suggests that repetitive DNA regions are particularly vulnerable to replication stress and DNA damage (Vona et al. 2018; Kreuz and Fischle 2016). We therefore lack a mechanistic framework connecting genotoxic insult, centromeric damage responses, and the cellular consequences of incomplete repair.

In disease contexts such as systemic sclerosis (SSc), a chronic autoimmune disorder characterized by fibrosis and immune dysregulation, yet the mechanistic links between genome instability and immune activation in SSc remain to be defined (Volkmann et al. 2023; Paul et al. 2022; van Bon et al. 2011; Rosendahl et al. 2022). A defining clinical feature of limited cutaneous SSc (lcSSc) is the presence of anti-centromere antibodies (ACAs), directed against centromeric proteins such as CENP-A and CENP-B (Kuwana et al. 1995; Akbarali et al. 2006). While recent studies have clarified immune pathways that sustain fibrosis (Brown and O’Reilly 2019; Wei et al. 2011; Ko et al. 2023; Usategui et al. 2011; Plikus et al. 2021), far less is known about how genome instability at repetitive DNA elements, including centromeres, contributes to chromosomal instability and centromeric antigen exposure, a process that may underlie the anti-centromere antibody response characteristic of lcSSc (Paul et al. 2022; Kirsch-Volders et al. 2020).

Damage within α-satellite DNA can disrupt kinetochore assembly and drive chromosomal instability (CIN), manifested by aneuploidy and micronucleus formation (Kreuz and Fischle 2016; Fachinetti et al. 2015; Krupina et al. 2021; Fenech et al. 2020). Mis-segregated chromosomes frequently form micronuclei that undergo rupture, releasing DNA into the cytoplasm. Cytosolic DNA can activate innate immune sensors such as cGAS, inducing interferon responses (Paul et al. 2022; Zhang et al. 2015; Mackenzie et al. 2017; Harding et al. 2017), and may enable interactions between nuclear-derived material and antigen-processing pathways (Gourh et al. 2020; Furukawa et al. 2016; Dengjel et al. 2005; Münz 2012). These observations suggest a potential link between repeat DNA instability and downstream cellular responses, although direct connections to centromere-derived antigen presentation remain to be established.

Bleomycin (BLM), a chemotherapeutic antibiotic and widely used experimental fibrosis inducer, generates reactive oxygen species that produce both single- and double-strand DNA breaks (DSBs) (Gülle et al. 2024; Chen and Stubbe 2005). Early studies demonstrated preferential BLM cleavage within repetitive DNA, including α-satellite sequences (Murray and Martin 1985; Hecht 2000; Chen et al. 2008; Murray et al. 2018). In vivo, BLM administration recapitulates key features of SSc, including dermal fibrosis, fibroblast activation, and extracellular matrix accumulation (Bi et al. 2023; Yoshizaki et al. 2010; Pawelec et al. 2022). These properties make BLM a useful model to examine how genotoxic stress affects centromeric DNA and genome stability in fibroblasts and *in vivo* (Paul et al. 2022; Waseem et al. 2025). However, the repair dynamics of centromeric DSBs lack mechanistic characterization. DSBs can be resolved by non-homologous end joining (NHEJ) or homologous recombination (HR), but in repetitive DNA, misalignment during HR can lead to deletions or insertions (Her and Bunting 2018; Aymard et al. 2014; Tsouroula et al. 2016). While CENP-A accumulation has been reported at induced damage sites (Zeitlin et al. 2009; De Rop et al. 2012), it is unclear whether this reflects recruitment of repair factors or the intrinsic fragility of centromeric chromatin. In our prior work, we demonstrated that SSc fibroblasts exhibit elevated centromeric DNA damage, micronucleus formation, and cGAS-STING activation (Paul et al. 2022); however, the mechanism of centromeric damage induction and the repair dynamics remained uncharacterized.

Here, we use BLM-induced DNA damage in fibroblasts and a mouse skin fibrosis model to investigate centromeric repeat instability and repair dynamics. Through α-satellite qPCR, immunofluorescence mapping, ATM inhibition, and patient fibroblast analyses, we demonstrate that BLM induces DNA double-strand breaks at active centromeres. These lesions are repaired primarily by RAD51-mediated homologous recombination but remain incompletely resolved, resulting in DNA loss, kinetochore disruption, missegregation, and micronucleus formation. Damaged centromere-associated chromatin appears in the cytoplasm following nuclear envelope perturbation, where it shows spatial proximity to HLA-DRB1. Together, these findings define a mechanistic framework linking active-centromere instability, ATM-dependent signaling, RAD51-mediated but error-prone repair, and downstream chromatin mislocalization.

## Results

### Genotoxic stress induces genome-wide centromeric repeat instability in vivo and in fibroblasts

To determine whether genotoxic stress induces instability across multiple centromeric loci genome-wide, we quantified α-satellite arrays representing centromeres from all human chromosomes in BLM-treated human fibroblasts and in the intradermal bleomycin mouse model. In human fibroblasts (CHON-002), BLM exposure (1.7–7.0 µM, 1–3 h) caused a rapid reduction in α-satellite DNA at 2 h, followed by partial recovery at 3 h, indicating dynamic, dose-dependent changes over time (Fig. 1F). At higher doses (7.0 µM), alterations persisted and failed to recover. After 24 h post-treatment recovery, copy number losses of up to ∼50% were sustained across chromosome-specific arrays including D1Z5, D6Z1, D15Z3, and D18Z1, demonstrating that centromere instability is not restricted to a single locus but affects centromeres genome-wide (Fig. 1G). BJ-5ta fibroblasts displayed a similar pattern but with greater severity, showing net losses and gains at comparable frequency and persistent losses at D5Z1, D7Z1, D9Z4, and D17Z1 (Supplementary Fig. 1A-B).

**Fig. 1.**
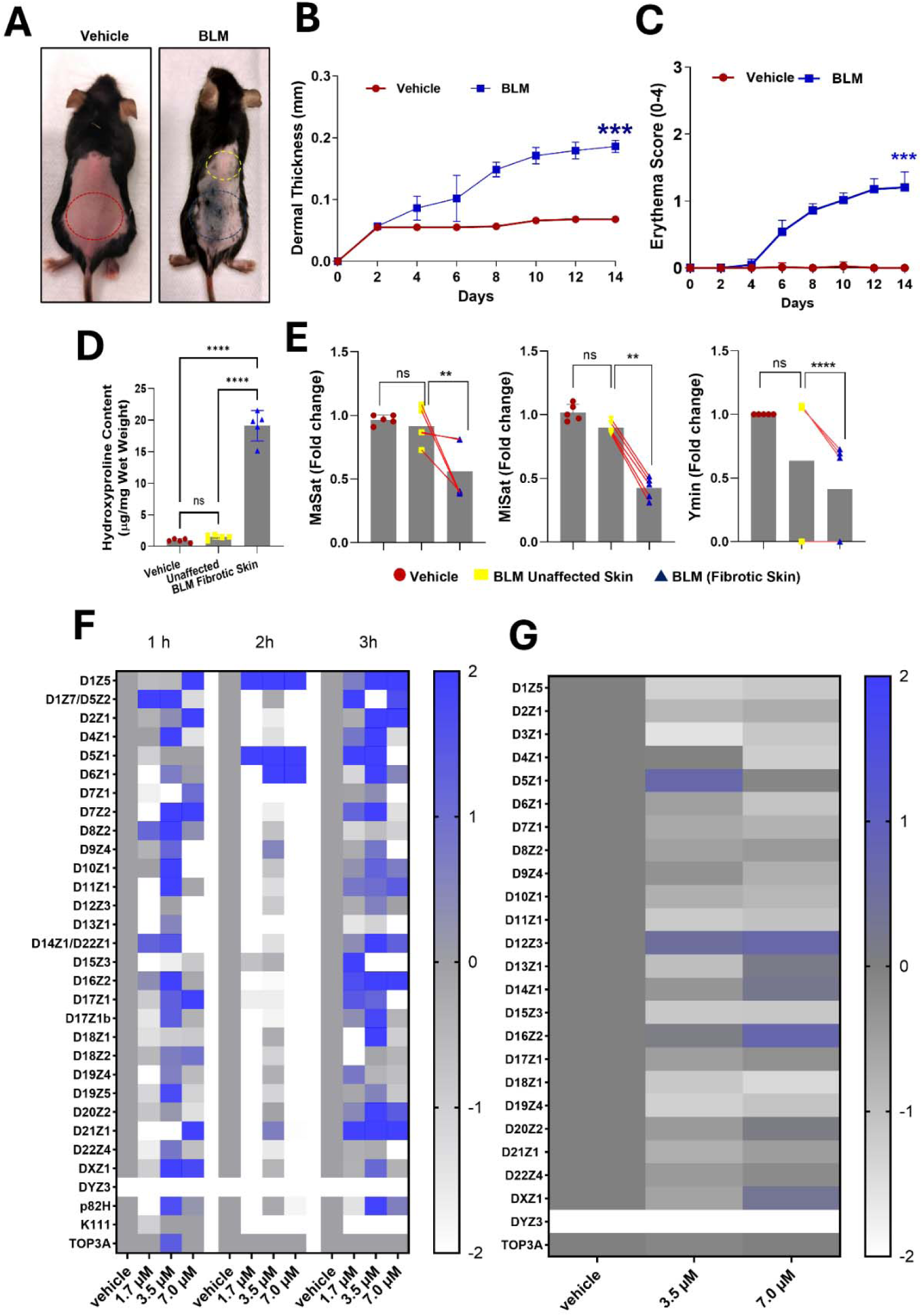
Bleomycin induces centromeric α-satellite alterations in mouse skin and human fibroblasts. (A) Representative dorsal skin images of mice after 14 days of vehicle (PBS) or bleomycin (BLM) treatment (n = 5 per group). Colors denote: vehicle control (red), adjacent unaffected skin (yellow), and fibrotic lesion (blue). (B-C) Quantification of dermal thickness (B) and erythema scores (C) in vehicle- and BLM-treated mice. (D) Hydroxyproline content measured by colorimetric assay in vehicle-control, adjacent unaffected, and fibrotic skin from BLM-treated mice. (E) qPCR analysis of mouse centromeric satellite repeats (MaSat, MiSat, Ymin) normalized to 18S. Red lines connect paired samples from adjacent unaffected and fibrotic skin within the same animal. (F-G) Heatmaps showing log fold change in human α-satellite monomers in CHON-002 fibroblasts following BLM treatment. (F) Cells treated with BLM for 1-3 h. (G) Cells treated with BLM for 3 h followed by 24 h recovery. Color scale: white, copy number loss; blue, copy number gain; grey, no change. DYZ3 was not detected in CHON-002 cells (female origin). TOP3A served as a non-centromeric reference. Data are presented as mean ± SD from three independent experiments. Statistical significance was determined by one-way ANOVA with appropriate post hoc testing: \*\**p* < 0.01; \*\*\**p* < 0.001; \*\*\*\**p* < 0.0001; ns, not significant.

MTT assays confirmed ∼75–80% viability at 3.5–7.0 µM (Supplementary Fig. 2), indicating that these copy number changes are not attributable to overt cytotoxicity.

To determine whether this instability extends to an in vivo fibrosis model, mice received intradermal BLM injections every other day for 14 days. By day 14, fibrotic skin showed marked increases in dermal thickness, erythema, and hydroxyproline content compared with vehicle controls (Fig. 1A-D), confirming successful fibrosis induction. qPCR of centromeric satellite repeats in fibrotic tissue revealed reductions of ∼50% for MaSat, ∼60% for MiSat, and ∼70% for Ymin relative to vehicle-injected skin (Fig. 1E), recapitulating the centromeric copy number losses observed in human fibroblasts and extending this finding to a disease-relevant in vivo context.

### Bleomycin induces double-strand breaks at active centromeres

We next examined whether BLM preferentially induces DSBs at centromeres marked by CENP-A. Here, we define active centromeres as CENP-A-containing chromatin domains, which mark functional kinetochore-associated centromeres. CHON-002 fibroblasts treated with 3.5 or 7.0 µM BLM for 3 h showed increased γH2AX signal compared with untreated controls, consistent with widespread DSB induction (Fig. 2A). Similar results were obtained in BJ-5ta fibroblasts (Supplementary Fig. 3B), and western blotting confirmed increased γH2AX levels following BLM exposure (Supplementary Fig. 3A).

**Fig. 2.**
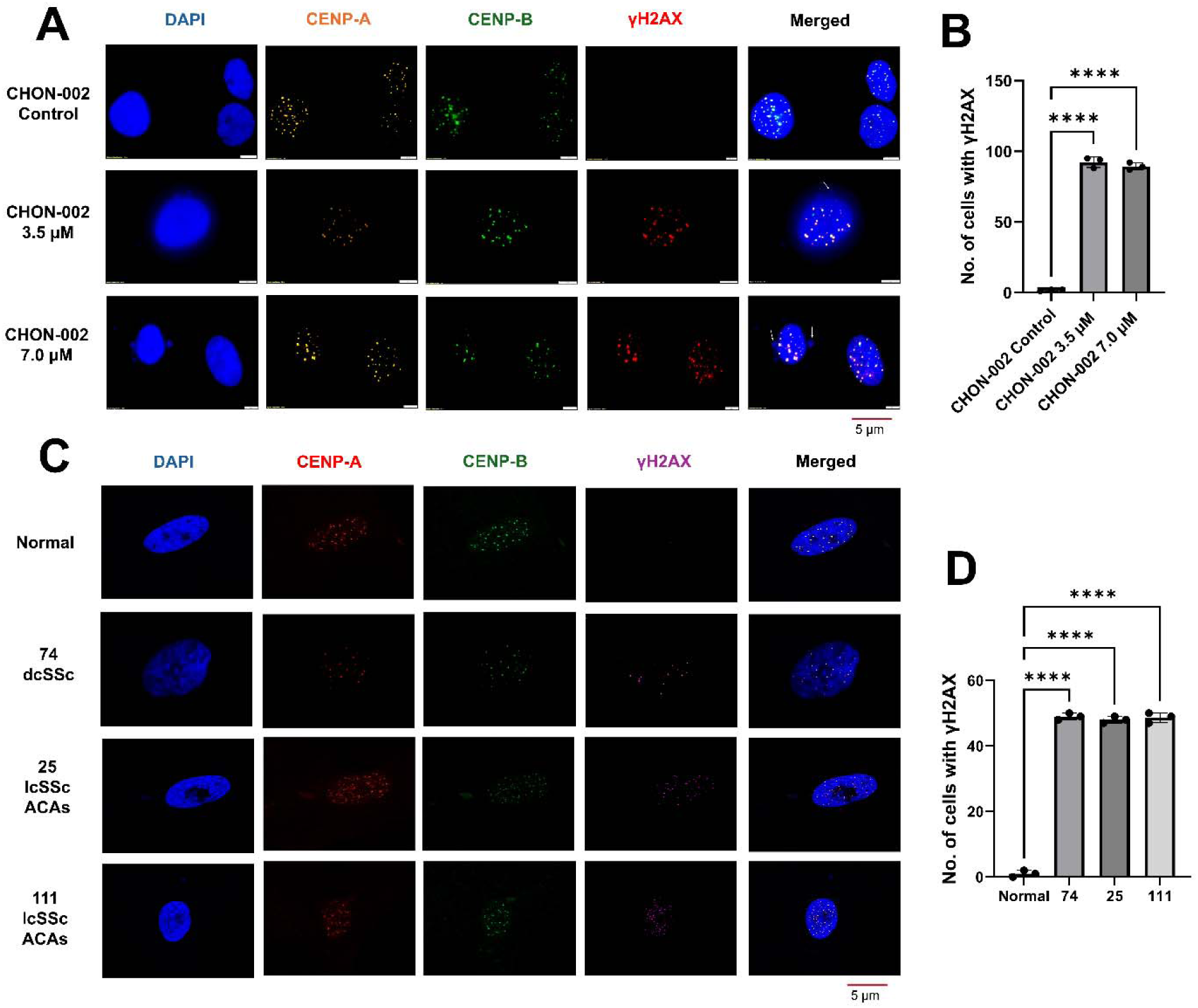
Genotoxic stress induces double-strand breaks at active centromeres. (A) Immunofluorescence analysis of CHON-002 fibroblasts treated with bleomycin (BLM; 3.5 or 7.0 µM, 3 h) and stained for CENP-A (orange), CENP-B (green), _γ_H2AX (red), and DAPI (blue, nuclei). White arrows indicate micronuclei. (B) Quantification of _γ_H2AX signal and association with centromeric markers (n = 100 cells). (C) Immunofluorescence analysis of primary dermal fibroblasts from healthy controls and SSc patients stained for CENP-A (green), _γ_H2AX (purple), and DAPI (blue). (D) Quantification of γH2AX fluorescence intensity (n = 50 cells). Data represent mean ± SD from three independent counts. Statistical significance was determined using unpaired t-test or one-way ANOVA: \**p* < 0.05; \*\**p* < 0.01; \*\*\**p* < 0.001; ns, not significant. Quantitative correlation analysis is shown in Fig. S3.

Pixel intensity correlation analysis revealed preferential association of γH2AX with active centromeric chromatin. While CENP-A and CENP-B signals remained tightly correlated (r > 0.85) under all conditions, γH2AX showed higher pixel intensity correlation with CENP-A (r = 0.51-0.74) than with CENP-B (r = 0.31–0.50), indicating that DSB signaling is enriched at CENP-A-marked centromeric domains rather than distributed across all α-satellite–associated chromatin (Supplementary Fig. 3C-D).

To determine whether centromeric DSB patterns are recapitulated in disease-relevant cells, we examined fibroblasts from three SSc patients as a preliminary dataset. In fibroblasts from two lcSSc patients, γH2AX showed centromeric association with CENP-A (Pearson’s r = 0.29-0.42) with markedly lower overlap with CENP-B (r = 0.09-0.19), mirroring the pattern observed in BLM-treated fibroblasts (Fig. 2C-D, Supplementary Fig. 3E). In contrast, fibroblasts from one dcSSc patient displayed γH2AX signal with minimal overlap with either centromeric marker (Fig. 2C, Supplementary Fig. 3E). Taken together, these findings demonstrate that lcSSc patient fibroblasts exhibit centromeric γH2AX association similar to BLM-treated cells, while the single dcSSc sample showed minimal overlap, a pattern consistent with the higher prevalence of anti-centromere antibodies in lcSSc. Given the limited sample size, these observations should be considered hypothesis-generating; definitive conclusions regarding disease subset-specific mechanisms will require larger patient cohorts.

### Centromeric breaks engage RAD51-mediated homologous recombination

To determine the DNA repair pathways engaged at centromeric DSBs, CHON-002 fibroblasts were treated with 3.5 or 7.0 µM BLM and immunostained for γH2AX (DSB marker), RAD51 (HR), and KU70 (NHEJ). Untreated cells showed minimal nuclear staining for all markers (Fig. 3A). BLM exposure induced γH2AX foci with spatially associated RAD51 signal, while KU70 showed markedly lower association with γH2AX, suggesting pathway-specific engagement at sites of damage.

**Fig. 3.**
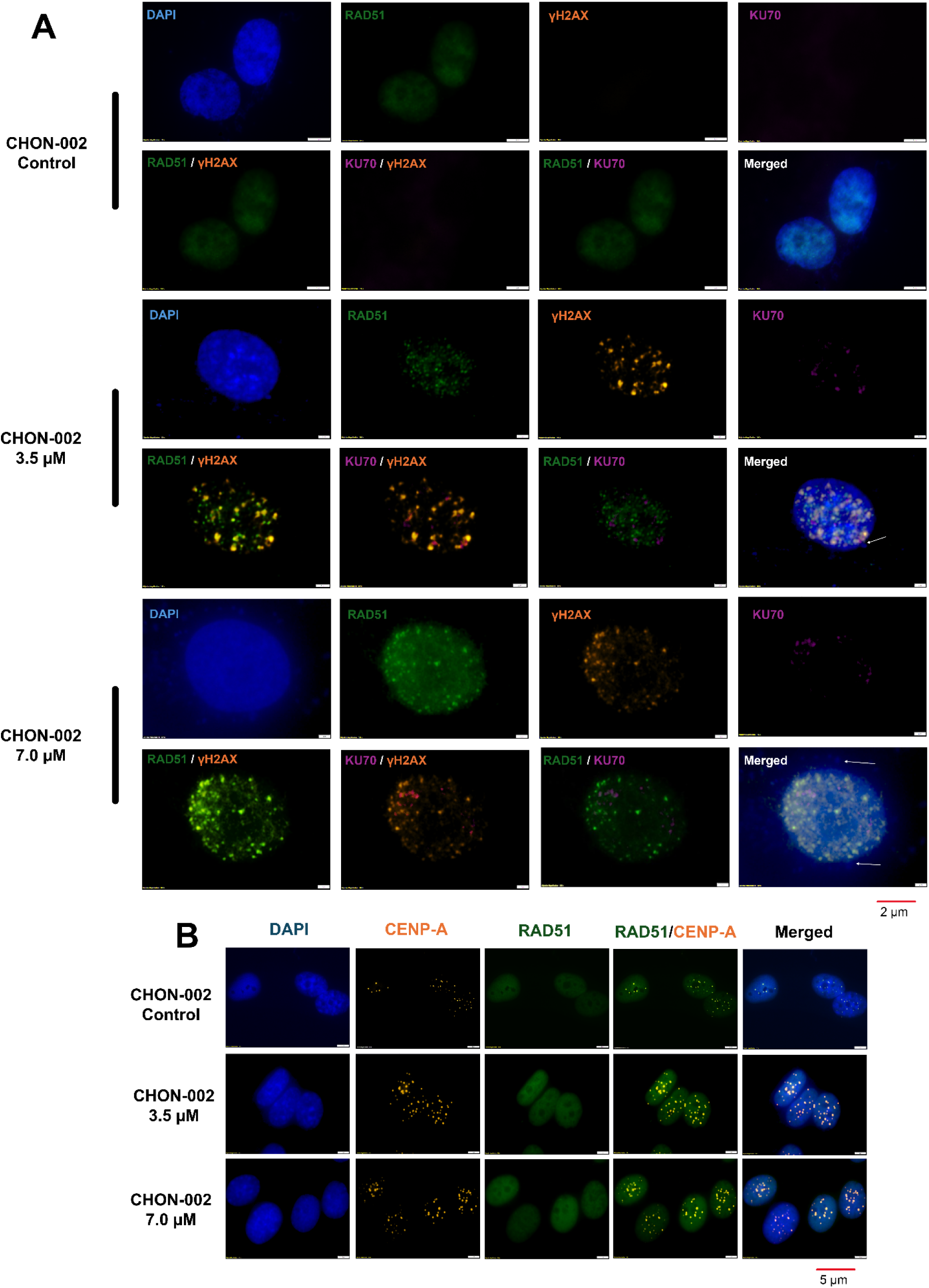
Centromeric breaks engage RAD51-mediated homologous recombination. CHON-002 fibroblasts were treated with bleomycin (BLM; 3.5 or 7.0 µM) for 3 h followed by 24 h recovery, unless otherwise indicated. (A) Immunofluorescence analysis of RAD51 (homologous recombination; green), γH2AX (DNA double-strand breaks; orange), KU70 (non-homologous end joining; purple), and DAPI (blue, nuclei). White arrows indicate micronuclei. (B) Immunofluorescence analysis of CENP-A (orange) and RAD51 (green) following 3 h BLM treatment (no recovery). RAD51 signal is shown in relation to centromeric regions marked by CENP-A. Quantitative correlation analysis of RAD51, KU70, and γH2AX signal association is shown in Fig. S4.

Pixel intensity correlation analysis confirmed the preferential recruitment of HR over NHEJ machinery at centromeric DSBs. RAD51-γH2AX spatial association was strong at 3.5 µM BLM (r = 0.875) and increased further at 7.0 µM (r = 0.919), whereas KU70-γH2AX association remained consistently low across both doses (r = 0.314 and r = 0.276, respectively) (Supplementary Fig. 4A). This differential was corroborated at the transcriptional level: RAD51 mRNA levels increased in a dose-dependent manner (*p* < 0.0001), while KU70 expression showed only a modest rise (*p* < 0.001), indicating that the HR pathway is both preferentially recruited and transcriptionally upregulated in response to BLM-induced centromeric damage (Supplementary Fig. 4B).

Kinetic analysis revealed a temporal progression in repair factor localization consistent with active HR engagement. At 3 h post-treatment, RAD51 staining was diffuse and pan-nuclear; by 24 h, discrete RAD51 foci had formed with spatial association to residual γH2AX signal, indicative of ongoing repair at persistent DSB sites (Fig. 3A-B). Together, these data demonstrate that centromeric DSBs induced by BLM are resolved predominantly through RAD51-mediated homologous recombination, with limited NHEJ involvement.

### ATM activity is required for **_γ_**H2AX and RAD51 recruitment at centromeric breaks

ATM kinase phosphorylates H2AX to generate γH2AX at sites of DNA damage. To test whether ATM mediates BLM-induced centromeric signaling, CHON-002 fibroblasts were treated with 3.5 or 7.0 µM BLM for 3 h, with or without the ATM inhibitor KU-55933 (35 µM, 1 h pretreatment). BLM induced dose-dependent increases in γH2AX signal with spatial association to the centromeric marker CENP-A (Fig. 4A). ATM inhibition substantially reduced γH2AX intensity at both doses while CENP-A localization remained intact, demonstrating that the damage signal, not the centromere itself, is ATM-dependent (Fig. 4A-C). ATM inhibition additionally suppressed RAD51 puncta accumulation (Fig. 4D), linking ATM activity directly to HR factor recruitment at centromeric damage sites.

**Fig. 4.**
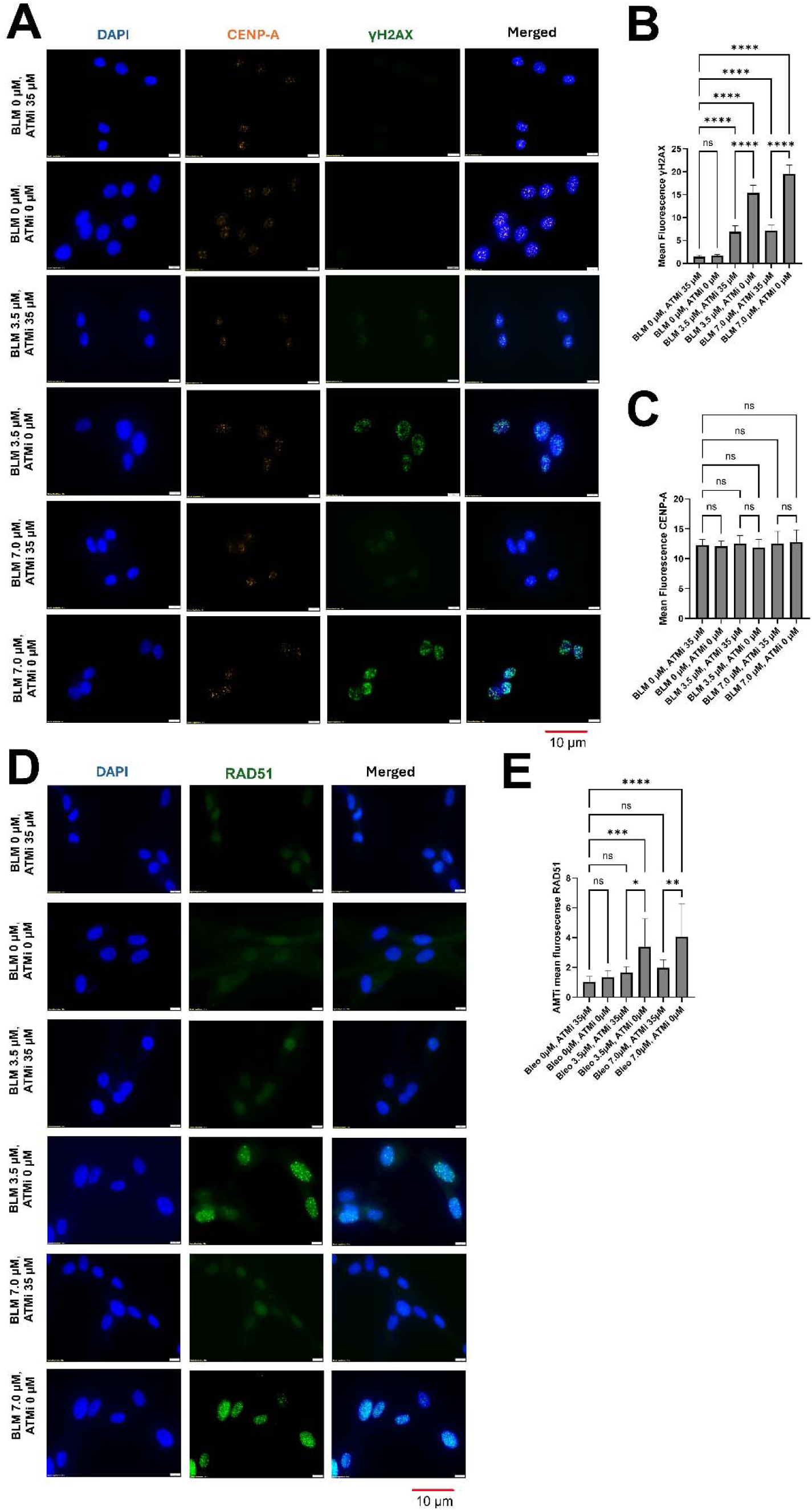
ATM activity is required for γH2AX and RAD51 at centromeres. CHON-002 fibroblasts were treated with bleomycin (BLM; 3.5 or 7.0 µM) for 3 h, with or without 1 h pretreatment with the ATM inhibitor KU-55933 (35 µM). (A) Immunofluorescence analysis of DAPI (blue, nuclei), CENP-A (orange, centromeres), and γH2AX (green, DNA double-strand breaks). γH2AX signal is shown in relation to centromeric regions marked by CENP-A. Scale bar, 10 µm. (B) Quantification of mean nuclear γH2AX intensity across conditions (n = 50 cells per condition). (C) Quantification of mean nuclear CENP-A intensity across conditions. (D) Immunofluorescence analysis of DAPI (blue, nuclei) and RAD51 (green, homologous recombination). Scale bar, 10 µm. (E) Quantification of mean nuclear RAD51 intensity across conditions (n = 50 cells per condition). Data are presented as mean ± SD. Statistical significance was determined by one-way ANOVA with appropriate post hoc testing: \**p* < 0.05; \*\**p* < 0.01; \*\*\**p* < 0.001; \*\*\*\**p* < 0.0001; ns, not significant.

Quantitative analysis confirmed that BLM-induced γH2AX and RAD51 induction were significantly suppressed by ATM inhibition (*p* < 0.001 and *p* < 0.0001, respectively; one-way ANOVA) (Fig. 4B, E), while CENP-A fluorescence levels were unaffected (Fig. 4C, ns). Together, these data demonstrate that ATM is required for both γH2AX induction and RAD51 recruitment at centromeric damage sites, placing centromeric DSB signaling within the canonical ATM-dependent DNA damage response.

### BLM-induced DNA damage activates PARP-dependent single-strand break signaling

In addition to DSBs, BLM generates single-strand breaks (SSBs) that activate PARP-dependent signaling. To determine whether this pathway is also engaged, we monitored poly(ADP-ribose) (PAR) polymer accumulation as a rapid marker of PARP activation. PAR immunofluorescence was minimal or absent in untreated CHON-002 fibroblasts (Supplementary Fig. 4C). Exposure to 7.0 µM BLM induced pan-nuclear PAR signal in ∼20% of cells, indicating PARP activation at sites of SSB damage; the remaining cells showed low or undetectable PAR signal, likely reflecting the well-established rapid turnover of PAR polymers following repair. Quantification confirmed a significant increase in nuclear PAR intensity relative to untreated controls (*p* < 0.0001, n = 10 cells per condition) (Supplementary Fig. 4D). These findings demonstrate that BLM engages both DSB-associated ATM–γH2AX signaling and SSB-associated PARP activation, indicating a broad genotoxic response at centromeric and flanking chromatin.

### Centromere damage promotes micronuclei formation and chromatin mislocalization

BLM treatment induced a significant increase in micronucleus frequency in both CHON-002 and BJ-5ta fibroblasts at 3.5 and 7.0 µM, whereas micronuclei were infrequent in untreated controls (Fig. 5A-B, Supplementary Fig. 5A, C). The majority of micronuclei contained centromeric signals, consistent with chromosome missegregation as the primary mechanism of formation and with prior observations in SSc fibroblasts (Paul et al. 2022). Critically, 30% of micronuclei contained CENP-B but lacked detectable CENP-A signal (Fig. 5A, C-D, Supplementary Fig. 5A-E), demonstrating loss of CENP-A-dependent centromere identity, consistent with eviction of the epigenetic mark that defines functional centromeres. This CENP-B/CENP-A signature represents a novel centromere-specific chromatin defect distinct from general chromosomal instability and suggests that genotoxic stress can uncouple the structural and functional components of centromere organization.

**Fig. 5.**
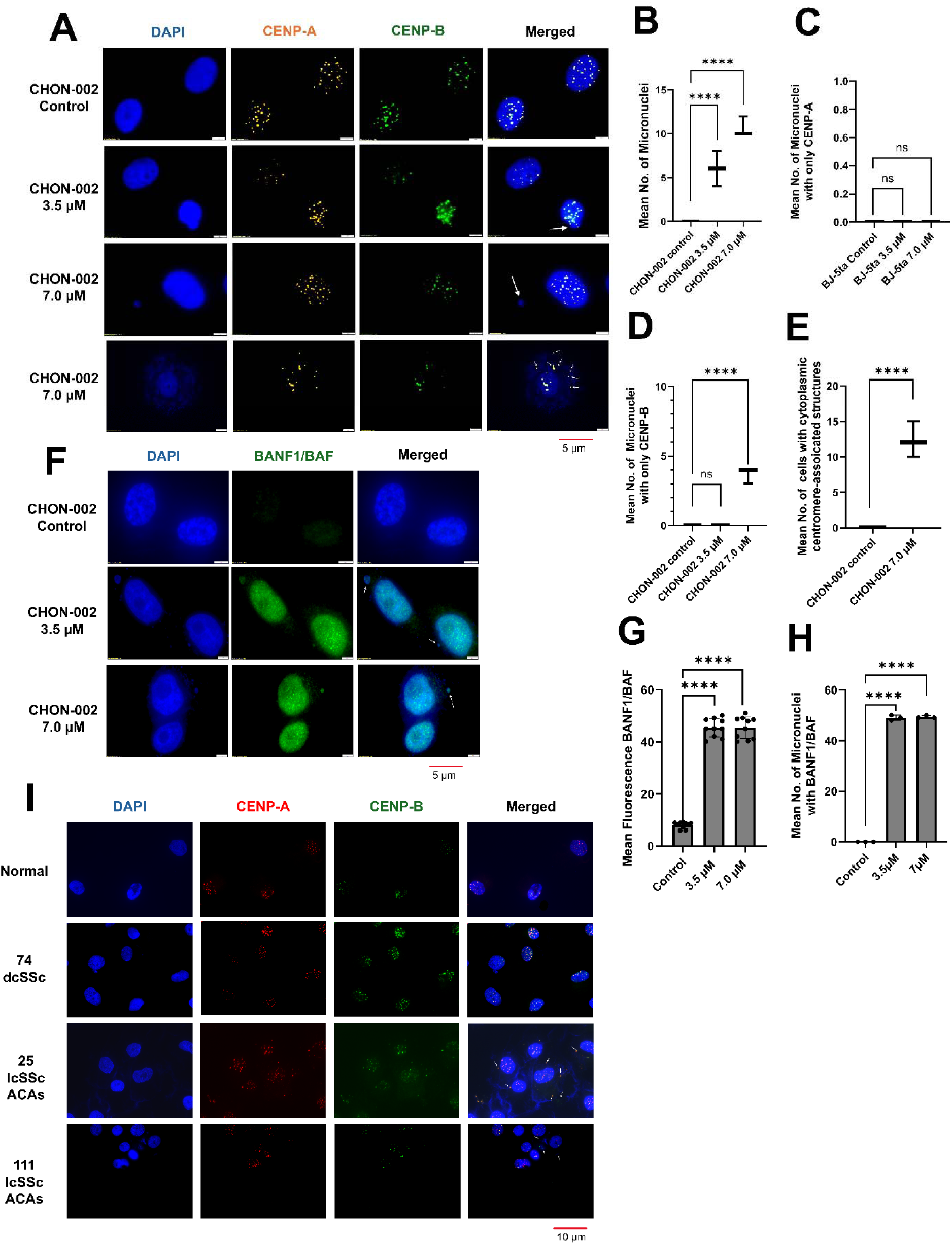
Centromere damage promotes micronuclei formation and chromatin mislocalization. CHON-002 fibroblasts were treated with bleomycin (BLM; 3.5 or 7.0 µM) for 3 h followed by 24 h recovery, unless otherwise indicated. (A) Immunofluorescence analysis of CENP-A (orange), CENP-B (green), and DAPI (blue, nuclei). White arrows indicate micronuclei; small white arrows indicate cytoplasmic centromere-associated structures. Approximately 30% of micronuclei shows CENP-B signal in the absence of detectable CENP-A, indicative of loss of CENP-A-dependent centromere identity. (B–E) Quantification of (B) mean number of micronuclei per 100 cells, (C) mean number of micronuclei positive for CENP-A, (D) mean number of micronuclei positive for CENP-B, and (E) mean number of cells containing cytoplasmic centromere-associated structures per 100 cells. At 7.0 µM BLM, ∼12% of CHON-002 cells displayed cytoplasmic centromere-associated structures (p < 0.0001; see also Supplementary Fig. 5F, where 15% of BJ-5ta fibroblasts showed the same phenotype under identical conditions). Quantifications are based on 100 cells per condition. Data represent mean ± SD from three independent experiments. (F) Immunofluorescence analysis of BANF1 (green) and DAPI (blue). BANF1 signal accumulates at rupture sites on primary nuclei and micronuclei in BLM-treated fibroblasts (white arrows), indicating nuclear envelope perturbation. (G) Quantification of mean BANF1 fluorescence intensity across conditions (n = 50 cells per condition). (H) Quantification of micronuclei positive for BANF1, expressed as the percentage of BANF1-positive micronuclei (n = 200 micronuclei). (I) Immunofluorescence analysis of primary dermal fibroblasts from healthy controls and systemic sclerosis (SSc) patients stained for CENP-A (red), CENP-B (green), and DAPI (blue). White arrows indicate cytoplasmic centromere-associated structures observed in two lcSSc patients (25 and 111) but not in a dcSSc patient (74). Data are presented as mean ± SD. Statistical significance was determined by one-way ANOVA with appropriate post hoc testing: ****p < 0.0001; ns, not significant.

A key consequence of centromere damage was the appearance of centromere-associated chromatin in the cytoplasm. At 7.0 µM BLM, ∼12% of CHON-002 cells displayed cytoplasmic centromere-associated structures, a significant and reproducible finding across three independent experiments (*p* < 0.0001; Fig. 5E). This was confirmed in BJ-5ta fibroblasts, where 15% of cells showed cytoplasmic centromere-associated structures under the same conditions (*p* < 0.0001, Supplementary Fig. 5F), establishing cytoplasmic mislocalization of centromeric chromatin as a consistent cellular response to BLM-induced genotoxic stress.

To determine the mechanism underlying this cytoplasmic release, we examined BANF1, a marker of nuclear envelope integrity that rapidly redistributes to sites of rupture. In untreated cells, BANF1 was confined to the nucleus; in BLM-treated fibroblasts, BANF1 strongly accumulated at rupture sites on both primary nuclei and micronuclei (white arrows, Fig. 5F). Quantification confirmed a significant increase in BANF1 intensity (*p* < 0.0001, n = 50 cells per condition; Fig. 5G) and a marked increase in the proportion of micronuclei positive for BANF1 (Fig. 5H), demonstrating that nuclear envelope rupture accompanies and likely mediates the cytoplasmic release of centromere-associated chromatin.

Primary fibroblasts from lcSSc patients recapitulated the cytoplasmic mislocalization phenotype. Fibroblasts from two ACA-positive lcSSc patients (patients 25 and 111) displayed frequent cytoplasmic centromere-associated structures, whereas fibroblasts from one dcSSc patient rarely did (Fig. 5I), a distribution consistent with the substantially higher prevalence of anti-centromere antibodies in lcSSc relative to dcSSc. Notably, cytoplasmic centromere-associated structures were previously identified in lcSSc fibroblasts (Paul et al. 2022), and the current findings extend this observation by demonstrating that equivalent structures can be experimentally induced through genotoxic stress and are mechanistically linked to nuclear envelope rupture. Together, the BANF1 redistribution data and the patient fibroblast findings establish that nuclear envelope rupture is a mechanistic conduit for centromeric chromatin mislocalization, and that this process is recapitulated in a disease subset where centromere-directed autoimmunity is a defining clinical feature.

### Cytoplasmic centromeric chromatin associates with MHC class II components

Having established that BLM-induced nuclear envelope rupture releases centromere-associated chromatin into the cytoplasm, we next asked whether this mislocalized material comes into contact with antigen-processing machinery. To address this, we performed immunofluorescence for CENP-B and HLA-DRB1, an MHC class II component expressed by fibroblasts in inflammatory contexts. In BLM-treated cells, we detected 10-15 discrete cytoplasmic CENP-B puncta per cell, whereas untreated cells lacked cytoplasmic CENP-B signal entirely. Object-based proximity analysis revealed that ∼90-95% of cytoplasmic CENP-B puncta were juxtaposed with punctate HLA-DRB1 signal (Fig. 6A). This association was observed exclusively in the ∼12% of cells displaying cytoplasmic centromere-associated structures and was absent in cells retaining intact nuclear CENP-B localization. DAPI signal was also detected within these cytoplasmic CENP-B-associated structures, consistent with the presence of chromatin-containing material. Untreated cells exhibited diffuse cytoplasmic HLA-DRB1 signal but lacked cytoplasmic CENP-B puncta, confirming that CENP-B–HLA-DRB1 juxtaposition is contingent on centromeric chromatin mislocalization rather than reflecting nonspecific HLA-DRB1 distribution.

**Fig. 6.**
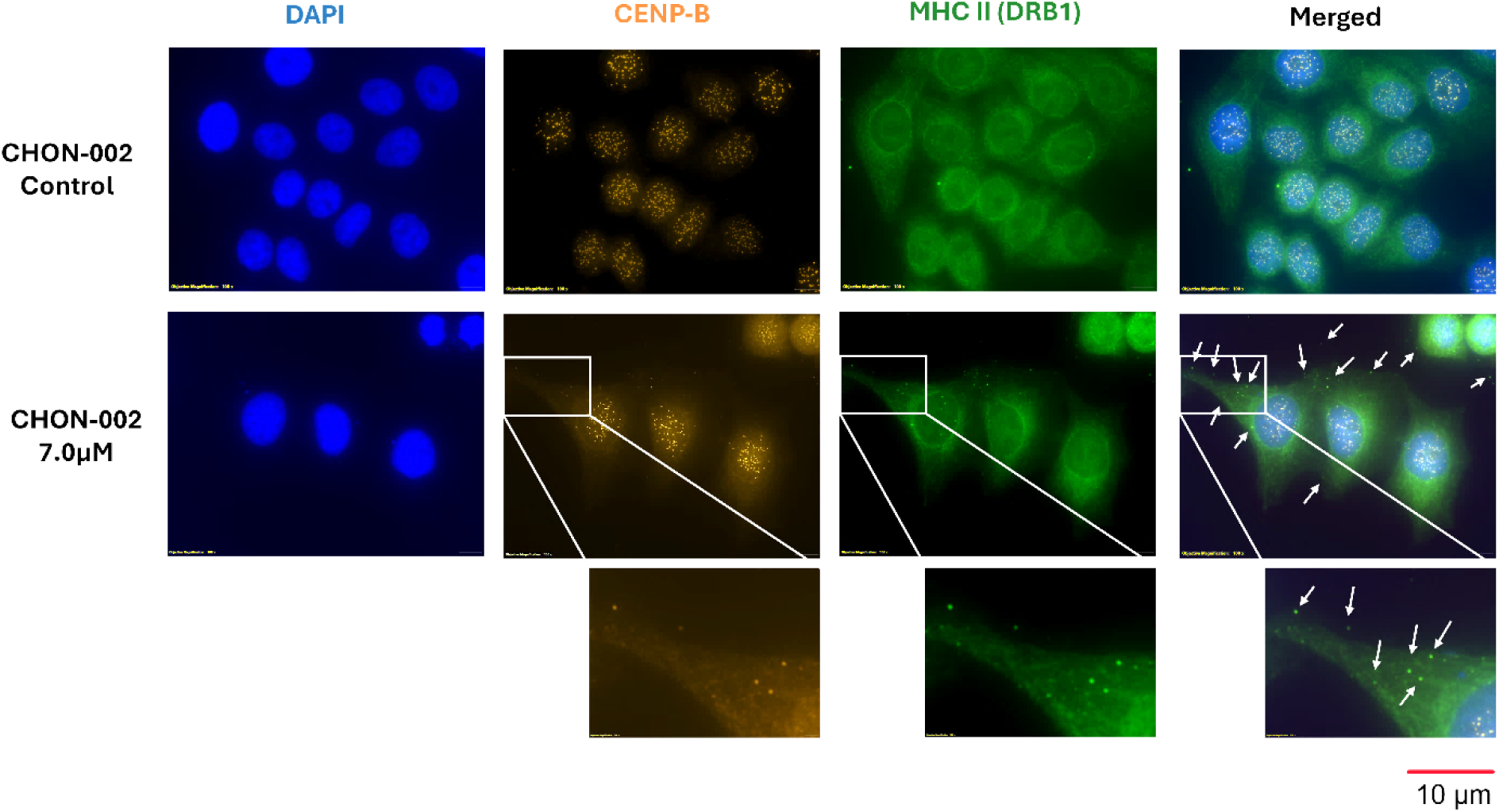
Association of cytoplasmic CENP-B with MHC class II (HLA-DRB1) in BLM-treated fibroblasts. CHON-002 fibroblasts were left untreated or treated with bleomycin (BLM; 7.0 µM) for 3 h followed by 24 h recovery. (A) Immunofluorescence analysis of CENP-B (orange), MHC class II (HLA-DRB1; green), and DAPI (blue, nuclei). White arrows indicat cytoplasmic CENP-B puncta juxtaposed with punctate HLA-DRB1 signal. Object-based proximity analysis showed that ∼90-100% of cytoplasmic CENP-B puncta (10-15 per cell) were associated with punctate HLA-DRB1 signal in BLM-treated cells. Untreated cells lacked cytoplasmic CENP-B puncta and served as specificity controls. The bottom panels show magnified views.

These findings should be interpreted as preliminary. Object-based proximity at confocal resolution does not establish direct molecular interaction, peptide processing, or antigen presentation. Nonetheless, the near-complete association of cytoplasmic CENP-B puncta with punctate HLA-DRB1 signal raises the possibility that nuclear envelope rupture creates a cytoplasmic compartment permissive for centromeric antigen encounter, a hypothesis that warrants direct functional testing in future studies. Notably, analogous proximity between cytoplasmic centromeric chromatin and MHC class II components has been reported in lcSSc fibroblasts (Paul et al. 2022), suggesting that this phenomenon is not limited to experimental genotoxic stress but may operate in disease-relevant contexts.

## Discussion

Centromeric α-satellite DNA has long been considered structurally protected by its specialized chromatin environment; our findings challenge this view by demonstrating that centromeres are not merely passive bystanders during genotoxic stress but are preferential targets of BLM-induced DNA damage. We show that BLM-induced breaks within centromeric α-satellite arrays are signaled through ATM-dependent γH2AX and repaired by RAD51-mediated homologous recombination, yet remain incompletely resolved, resulting in net copy number losses and gains across multiple chromosome-specific arrays. These findings position centromeres as active participants in genome stress responses rather than solely structural elements, and demonstrate that centromeric instability occurs genome-wide, reflecting a broad susceptibility of α-satellite repeats to genotoxic insult.

Centromeric repeats are intrinsically vulnerable due to their repetitive sequence composition, replication challenges, and specialized chromatin organization. Prior studies have shown that centromeres accumulate spontaneous lesions during both proliferation and quiescence, with repair occurring predominantly through RAD51-mediated HR (Nassar et al. 2023; Black and Giunta 2018; Saayman et al. 2023). In this study, BLM exposure amplified this baseline fragility, generating losses and gains within α-satellite arrays whose magnitude differed between fibroblast lines, suggesting that chromatin state or repair capacity modulates centromeric vulnerability. The BLM-induced skin fibrosis model recapitulated these reductions in centromeric satellite content in vivo (Chen and Stubbe 2005; Murray and Martin 1985; Hecht 2000; Chen et al. 2008), supporting centromeric DNA as a sensitive substrate for genotoxic stress across experimental systems.

Spatial association of γH2AX with CENP-A, and suppression of this signal by ATM inhibition, demonstrate that centromeric lesions engage canonical ATM-dependent DNA damage signaling. Preferential recruitment of RAD51 with limited KU70 involvement indicates that HR is the dominant repair pathway at these sites. However, HR did not fully restore centromeric array integrity, in line with prior observations that repair within repetitive DNA can be error-prone due to template misalignment or replication-associated stress (Black and Giunta 2018; Saayman et al. 2023; Mitra et al. 2014).

A particularly notable finding of this study is the identification of micronuclei retaining CENP-B but lacking detectable CENP-A, a CENP-B/CENP-A signature that has not been previously described as a consequence of acute genotoxic stress. CENP-A is the histone H3 variant that epigenetically defines centromere identity and is essential for kinetochore assembly and faithful chromosome segregation; its loss from a chromatin domain effectively renders that domain non-functional as a centromere, regardless of the underlying α-satellite sequence (Jeffery et al. 2022). In contrast, CENP-B binds constitutively to CENP-B boxes within α-satellite DNA independently of centromere activity and is retained even at inactive centromeres. The CENP-B/CENP-A configuration observed in ∼30% of micronuclei therefore indicates that genotoxic stress can uncouple the structural and epigenetic components of centromere organization, a defect mechanistically distinct from general chromosomal instability. Whether this reflects incomplete CENP-A replenishment following repair, active eviction of CENP-A from damaged chromatin, or preferential incorporation of canonical H3 at repair sites remains to be determined. These possibilities are not mutually exclusive and may reflect the known competition between CENP-A and H3.3 at centromeric chromatin during DNA damage responses (Jeffery et al. 2022). Regardless of mechanism, the loss of CENP-A from micronuclear centromeres suggests that even if the underlying DNA sequence is partially preserved following incomplete HR, the epigenetic information required to reconstitute a functional centromere may not be.

Functionally, centromere instability was associated with chromosome missegregation, micronucleus formation, and nuclear envelope rupture marked by BANF1 redistribution and cytoplasmic appearance of centromere-associated chromatin. These phenotypes were consistent with features previously described in SSc fibroblasts, particularly in limited cutaneous SSc (Paul et al. 2022). The observed juxtaposition of cytoplasmic CENP-B with HLA-DRB1 raises the possibility that mislocalized centromeric chromatin may come into contact with antigen-processing compartments, consistent with prior work linking cytosolic DNA to innate immune signaling pathways including cGAS-STING (Paul et al. 2022; Denais et al. 2016; Raab et al. 2016; Kono et al. 2022; Guey et al. 2020) and with the well-established association between anti-centromere antibodies and lcSSc (Kuwana et al. 1995).

Differences observed between fibroblast lines, including lcSSc and dcSSc samples, suggest variability in centromere damage responses and repair dynamics. However, the limited number of patient-derived lines analyzed here precludes definitive conclusions regarding disease subset-specific mechanisms.

This study has limitations. Fibroblast cultures and the BLM model do not fully capture the complexity of stromal-immune interactions in vivo, and direct demonstration of centromeric peptide processing, antigen presentation, and downstream immune activation remains to be established. Pixel intensity correlation analyses reflect physical juxtaposition at the resolution of confocal microscopy and do not establish direct molecular interactions or antigen presentation. In summary, these findings define a mechanistic framework linking active-centromere instability, ATM-dependent signaling, RAD51-mediated but error-prone repair, and downstream chromatin mislocalization in fibroblasts. Our findings provide a tractable experimental platform to dissect how centromeric DNA damage drives chromosomal instability and potentially seeds cytoplasmic DNA sensing pathways implicated in SSc pathogenesis. Future work should test whether pharmacological reinforcement of centromeric repair, for example, through modulation of ATM activity or RAD51 loading, can reduce micronucleus formation, limit nuclear envelope rupture, and attenuate downstream immune activation in disease-relevant models

## Methods

### Reagents

Bleomycin sulfate (Cayman Chemical, Cat. No. 13877, Lot #0701763-49) was used in all experiments. Reagent lot information is provided to ensure experimental reproducibility. All other reagents were of analytical grade and obtained from commercial suppliers unless otherwise noted.

### Cell culture and treatments

hTERT-immortalized human fibroblasts (CHON-002, ATCC CRL-2847; BJ-5ta, ATCC CRL-4001) were cultured in Dulbecco’s Modified Eagle Medium (DMEM; Gibco) supplemented with 10% fetal bovine serum (FBS; Gibco) and 1% penicillin-streptomycin (Gibco). Cells were maintained at 37 °C in a humidified incubator with 5% CO and routinely tested for mycoplasma contamination.

For acute bleomycin (BLM) exposure, fibroblasts were seeded at 2-3 × 10 cells per well in 6-well plates and grown to 60-80% confluence. Cells were treated with 0, 3.5, or 7 µM BLM for 3 h. For recovery assays, cells were washed with phosphate-buffered saline (PBS) and incubated in drug-free medium for 24 h. For ATM inhibition, cells were pretreated with KU-55933 (35 µM, Selleck Cat. S1092) for 1 h prior to BLM treatment.

### Mouse BLM-induced skin fibrosis model

C57BL/6 mice (Jackson Laboratory; 20-25 g; 2 males, 3 females per group) were randomized to receive intradermal injections of BLM (1 U/mL, 100 µL) or vehicle (PBS) into shaved dorsal skin every other day for 14 days. On day 14, mice were anesthetized with isoflurane (1-2%) and euthanized by cervical dislocation. Skin samples were collected from: (1) PBS-injected control sites, (2) adjacent uninjected skin from the same animal, and (3) fibrotic BLM-injected lesions. Genomic DNA was isolated for qPCR analysis. All animal studies were approved by the University of Alabama at Birmingham Institutional Animal Care and Use Committee (protocol 23172).

### Human samples

Primary dermal fibroblasts were derived from patients with SSc under University of Michigan IRB HUM00065044, as previously described (Paul et al. 2022). Cell lines were obtained from patients with limited cutaneous SSc (lcSSc; 25 and 111) and diffuse cutaneous SSc (dcSSc; 74), as well as from healthy controls. No identifiable patient information was used in this study.

### DNA and RNA isolation

Genomic DNA was extracted using the Monarch Genomic DNA Purification Kit (NEB 30102) with RNase A treatment. Total RNA was isolated using the Direct-zol RNA MicroPrep Kit (Zymo Research) with on-column DNase digestion. Concentrations were measured with a Qubit 4 Fluorometer (Thermo Fisher). DNA and RNA were stored at −80 °C until use.

### Centromere qPCR

Mouse centromeric arrays (MaSat, MiSat, and Ymin) were quantified by qPCR using validated primer sets (Benedetti et al. 2024). Human α-satellite arrays (D1-D22, X, Y) were quantified using primer sets described previously (Contreras-Galindo et al. 2017). Reactions were run on QuantStudio 3 (Applied Biosystems) with Radiant SYBR Green Master Mix (Alkali Scientific). Relative copy number was normalized to TOP3A (human) or 18S (mouse), unless otherwise indicated. Primer specificity was confirmed by melt-curve analysis. All reactions were performed in technical duplicates unless otherwise indicated.

### Western blotting

Cells were lysed in RIPA buffer (ChemCruz) supplemented with protease and phosphatase inhibitors (Thermo Fisher). Lysates (20-30 µg protein) were resolved by SDS-PAGE (10%) and transferred to PVDF membranes (Bio-Rad). Membranes were blocked in 5% milk/TBST, incubated overnight at 4 °C with primary antibodies (Supplementary Table 1), and probed with HRP-conjugated secondary antibodies (2 h, room temperature). Signals were developed using Clarity ECL substrate (Bio-Rad) and imaged with a ChemiDoc MP system. Band intensities were quantified using Image Lab (Bio-Rad). All quantifications were performed using Image Lab (Bio-Rad) with identical exposure conditions across samples.

### Immunofluorescence microscopy

Cells were fixed in 4% paraformaldehyde (PFA), permeabilized with PBST (0.2% Triton X-100), and blocked in 2% BSA/PBST. Coverslips were incubated with primary antibodies (Supplementary Table 1) for 1 h at 37 °C, followed by Alexa Fluor-conjugated secondary antibodies (1:1000, 45 min, room temperature). Nuclei were counterstained with DAPI (ProLong Gold, Thermo Fisher).

For chromosome spreads, cells were arrested with 10 µg/mL colcemid for 16 h, incubated in 0.075 M KCl for 15 min, cytospun at 1500 rpm (∼300 × g; Cytospin 3, Thermo Scientific) for 5 min, and fixed in 4% PFA prior to IF staining. Images were acquired on an Olympus BX73 microscope using identical exposure settings across conditions and processed uniformly using Fiji/ImageJ. Pixel intensity correlations (Pearson’s r) were calculated using the Coloc2 plugin in Fiji/ImageJ with thresholds set automatically using the Costes method. No post-acquisition intensity adjustments were applied differentially between conditions.

For CENP-B and HLA-DRB1 proximity analysis, cytoplasmic CENP-B puncta were identified manually in BLM-treated cells based on discrete punctate signal absent from untreated controls. For each punctum, HLA-DRB1 fluorescence intensity was assessed within the same cytoplasmic region. Untreated cells, which lacked cytoplasmic CENP-B puncta, served as specificity controls.

### Quantitative RT-PCR

mRNA expression of RAD51 and KU70 was measured using the Luna One-Step RT-qPCR Kit (NEB) with 50 ng RNA per reaction on a QuantStudio 3 instrument. GAPDH was used as a reference gene, and relative expression was calculated using the ΔΔCt method. Reactions were performed in technical duplicates. Primer sequences were obtained from published sources (Wang et al. 2015; Lim et al. 2002).

### Statistical analysis

All qPCR experiments were performed in three independent biological replicates with technical duplicates. Immunofluorescence quantifications were performed on at least 50-100 cells per condition from randomly selected fields, unless otherwise specified. Western blotting was performed in at least three independent replicates. Statistical significance was determined by unpaired two-tailed Student’s t-test or one-way ANOVA with Tukey’s or Dunnett’s post hoc test, as appropriate (GraphPad Prism v9.3.1). A threshold of p < 0.05 was considered significant. Pearson’s correlation coefficients (r) were calculated from pixel intensity correlations using the Coloc2 plugin in Fiji/ImageJ. Thresholds were set automatically using the Costes method and applied consistently across all conditions.

## Data Access

This study did not generate sequencing, array, or other large-scale genomics datasets requiring deposition in GEO, SRA, or BioProject. All source data supporting the findings of this study are provided with the manuscript. Additional materials are available from the corresponding author upon reasonable request.

## Acknowledgments

We thank Dr. Arko Sen for insightful discussions. We thank the National Scleroderma Foundation for supporting this project. R.C.-G. is the recipient of the Marta Marx Award for the Eradication of Scleroderma from the National Scleroderma Foundation.

## Author Contributions

R.C-G. and A.I. conceived and designed the study. R.C-G., A.I., M.W. and C.W performed the investigation. R.C-G acquired the funding. R.C-G., M.W., B.C., and W.C supervised the study. R.C-G., M.W., H.O., B.C., and W.C. wrote, reviewed, and edited the manuscript.

## References

Akbarali Y, Ogasawara H, Hasegawa M, Fujimoto M, Matsushita T, Hamaguchi Y, Komura K, Takehara K, Sato S. 2006. Fine specificity mapping of autoantigens targeted by anti-centromere autoantibodies. J Autoimmun 27: 272–280.

Aldrup-Macdonald ME, Sullivan BA. 2014. The past, present, and future of human centromere genomics. Genes 5: 33–50.

Aymard F, Bugler B, Schmidt CK, Guillou E, Caron P, Briois S, Iacovoni JS, Daburon V, Miller KM, Jackson SP, et al. 2014. Transcriptionally active chromatin recruits homologous recombination at DNA double-strand breaks. Nat Struct Mol Biol 21: 366–374.

Benedetti F, Curreli S, Gualtieri A, Obino V, Cotello S, Caputo A, Nigro A, Peruzzi F, Schifano F, Khalili K, et al. 2024. Mycoplasma DnaK expression increases cancer development in vivo upon DNA damage. Proc Natl Acad Sci USA 121: e2320859121.

Bi X, Mills T, Wu M. 2023. Animal models in systemic sclerosis: an update. Curr Opin Rheumatol 35: 364–370.

Black EM, Giunta S. 2018. Repetitive fragile sites: centromere satellite DNA as a source of genome instability in human diseases. Genes 9: 615.

Brown M, O’Reilly S. 2019. The immunopathogenesis of fibrosis in systemic sclerosis. Clin Exp Immunol 195: 310–321.

Chen J, Stubbe JA. 2005. Bleomycin: toward better therapeutics. Nat Rev Cancer 5: 102–112.

Chen J, Ghorai MK, Kenney G, Stubbe J. 2008. Mechanistic studies on bleomycin-mediated DNA damage: multiple binding modes can result in double-stranded DNA cleavage. Nucleic Acids Res 36: 3781–3790.

Contreras-Galindo R, Fischer S, Saha A, Wainberg M, Celi F, McDonald JF, Brown TA. 2017. Rapid molecular assays to study human centromere genomics. Genome Res 27: 2040–2049.

De Rop V, Padeganeh A, Maddox PS. 2012. CENP-A: the key player behind centromere identity, propagation, and kinetochore assembly. Chromosoma 121: 527–538.

Denais CM, Gilbert RM, Isermann P, McGregor AL, te Lindert M, Weigelin B, Davidson PM, Friedl P, Wolf K, Lammerding J. 2016. Nuclear envelope rupture and repair during cancer cell migration. Science 352: 353–358.

Dengjel J, Schoor O, Fischer R, Reich M, Kraus M, Muller M, Kreymborg K, Altenberend F, Brandenburg J, Kalbacher H, et al. 2005. Autophagy promotes MHC class II presentation of peptides from intracellular source proteins. Proc Natl Acad Sci USA 102: 7922–7927.

Fachinetti D, Han JS, McMahon MA, Ly P, Abdullah A, Wong AJ, Cleveland DW. 2015. DNA sequence-specific binding of CENP-B enhances the fidelity of human centromere function. Dev Cell 33: 314–327.

Fenech M, Knasmueller S, Bolognesi C, Bonassi S, Holland N, Migliore L, Palitti F, Natarajan AT, Kirsch-Volders M. 2020. Micronuclei as biomarkers of DNA damage, aneuploidy inducers of chromosomal hypermutation, and as sources of pro-inflammatory DNA in humans. Mutat Res Rev Mutat Res 786: 108342.

Furukawa H, Oka S, Shimada K, Sugii S, Ohashi J, Matsui T, Ikenaka T, Nakayama H, Hashimoto A, Takaoka H, et al. 2016. Human leukocyte antigen and systemic sclerosis in Japanese: the sign of the four independent protective alleles. PLoS One 11: e0154255.

Gourh P, Safran SA, Alexander T, Boyden SE, Morgan ND, Shah AA, Criswell LA, Stevens AM, Mayes MD, Arnett FC, et al. 2020. HLA and autoantibodies define scleroderma subtypes and risk in African and European Americans and suggest a role for molecular mimicry. Proc Natl Acad Sci USA 117: 552–562.

Guey B, Wischnewski M, Decout A, Makasheva K, Kaynak M, Sakar MS, Fierz B, Ablasser A. 2020. BAF restricts cGAS on nuclear DNA to prevent innate immune activation. Science 369: 823–828.

Gülle S, Gecit I, Dumankaya S, Kargili A, Yildirim A, Yayla S, Yildiz F. 2024. Skin and lung fibrosis induced by bleomycin in mice: a systematic review. Reumatismo 76: 22–32.

Harding SM, Benci JL, Irianto J, Discher DE, Minn AJ, Greenberg RA. 2017. Mitotic progression following DNA damage enables pattern recognition within micronuclei. Nature 548: 466–470.

Hecht SM. 2000. Bleomycin: new perspectives on the mechanism of action. J Nat Prod 63: 158–168.

Her J, Bunting SF. 2018. How cells ensure correct repair of DNA double-strand breaks. J Biol Chem 293: 10502–10511.

Jeffery D, Lochhead M, Almouzni G. 2022. CENP-A: a histone H3 variant with key roles in centromere architecture in healthy and diseased states. Results Probl Cell Differ 70: 221–261.

Kirsch-Volders M, Bolognesi C, Ceppi M, Bruzzone M, Fenech M. 2020. Micronuclei, inflammation, and autoimmune disease. Mutat Res Rev Mutat Res 786: 108335.

Ko J, Ahn J, Kim J, Park D, Lee SH. 2023. The pathogenesis of systemic sclerosis: the origin of fibrosis and interlink with vasculopathy and autoimmunity. Int J Mol Sci 24: 14287.

Kono Y, Kubota T, Tamura M, Aoki K, Hieda M, Kamura T, Haraguchi T, Hiraoka Y, Kimura H, Shimi T. 2022. Nucleoplasmic lamin C rapidly accumulates at sites of nuclear envelope rupture with BAF and cGAS. J Cell Biol 221: e202201024.

Kreuz S, Fischle W. 2016. Oxidative stress signaling to chromatin in health and disease. Epigenomics 8: 843–862.

Krupina K, Goginashvili A, Cleveland DW. 2021. Causes and consequences of micronuclei. Curr Opin Cell Biol 70: 91–99.

Kuwana M, Okano Y, Pandey JP, Silver RM, Fertig N, Medsger TA Jr, Wright TM. 1995. HLA class II genes associated with anticentromere antibody in systemic sclerosis (scleroderma). Ann Rheum Dis 54: 983–987.

Lim JW, Kim H, Kim KH. 2002. Expression of Ku70 and Ku80 mediated by NF-κB and cyclooxygenase-2 is related to proliferation of human gastric cancer cells. J Biol Chem 277: 46093–46100.

Logsdon GA, Vollger MR, Hsieh P, Mao Y, Liskovykh M, Koren S, Nurk S, Mercuri L, Dishuck PC, Rhie A, et al. 2025. Complex genetic variation in nearly complete human genomes. Nature 644: 430–441.

Mackenzie KJ, Carroll P, Martin CA, Murina O, Fluteau A, Simpson DJ, Olova N, Sutcliffe H, Rainger JK, Leitch A, et al. 2017. cGAS surveillance of micronuclei links genome instability to innate immunity. Nature 548: 461–465.

McNulty SM, Sullivan BA. 2018. Alpha-satellite DNA biology: finding function in the recesses of the genome. Chromosome Res 26: 115–138.

Mitra S, Gomez-Raja J, Larriba G, Dubey DD, Sanyal K. 2014. Rad51–Rad52-mediated maintenance of centromeric chromatin in Candida albicans. PLoS Genet 10: e1004344.

Münz C. 2012. Antigen processing for MHC class II presentation via autophagy. Front Immunol 3: 9.

Murray V, Martin RF. 1985. The sequence specificity of bleomycin-induced DNA damage in intact cells. J Biol Chem 260: 10389–10391.

Murray V, Chen JK, Chung LH. 2018. The interaction of the metallo-glycopeptide anti-tumour drug bleomycin with DNA. Int J Mol Sci 19: 1372.

Nassar R, Thompson L, Fouquerel E. 2023. Molecular mechanisms protecting centromeres from self-sabotage and implications for cancer therapy. NAR Cancer 5: zcad019.

Paul S, Imtiaz A, Wang S, Lee H, Vaidya A, Chatterjee S, Czaja W, Contreras-Galindo R. 2022. Centromere defects, chromosome instability, and cGAS-STING activation in systemic sclerosis. Nat Commun 13: 7074.

Pawelec KM, Tchao J, Dasgupta A, Fan Y, Clauss M, Chandel NS, Budinger GRS, Karmouty-Quintana H, McVerry BJ, Ridge KM. 2022. Prevention of bleomycin-induced lung fibrosis via inhibition of the MRTF/SRF transcription pathway. Pharmacol Res Perspect 10: e01028.

Plikus MV, Wang X, Sinha S, Forte E, Thompson SM, Herzog EL, Driskell RR, Rosenthal N, Biernaskie J, Horsley V. 2021. Fibroblasts: origins, definitions, and functions in health and disease. Cell 184: 3852–3872.

Raab M, Gentili M, de Belly H, Thiam HR, Vargas P, Jimenez AJ, Lautenschlaeger F, Voituriez R, Lennon-Dumenil AM, Manel N, et al. 2016. ESCRT III repairs nuclear envelope ruptures during cell migration to limit DNA damage and cell death. Science 352: 359–362.

Rosendahl AH, Schönborn K, Krieg T. 2022. Pathophysiology of systemic sclerosis (scleroderma). Kaohsiung J Med Sci 38: 187–195.

Saayman X, Hayward A, Collins JA, Titus KR, Chotalia M, Dagg RA, Grubb C, Pontano LL, Bonilla B, McKinnon PJ, et al. 2023. Centromeres as universal hotspots of DNA breakage driving RAD51-mediated recombination during quiescence. Mol Cell 83: 523–538.e7.

Savigny F, Schricke C, Lecerf C, Deslée G, Césaire T, Pinet C, Dubucquoi S, Sokol H, Crestani B, Mailleux AA, et al. 2021. Protective role of the nucleic acid sensor STING in pulmonary fibrosis. Front Immunol 11: 588799.

Tsouroula K, Furst A, Rogier M, Heyer V, Maglott-Roth A, Ferrand A, Reina-San-Martin B, Soutoglou E. 2016. Temporal and spatial uncoupling of DSB repair pathways within mammalian heterochromatin. Mol Cell 63: 293–305.

Usategui A, del Rey MJ, Pablos JL. 2011. Fibroblast abnormalities in the pathogenesis of systemic sclerosis. Expert Rev Clin Immunol 7: 491–498.

van Bon L, Cossu M, Radstake TR. 2011. An update on an immune system that goes awry in systemic sclerosis. Curr Opin Rheumatol 23: 505–510.

Volkmann ER, Andréasson K, Smith V. 2023. Systemic sclerosis. Lancet 401: 304–318.

Vona R, Iannace A, Gambardella L, Nocella C, Cammisotto V, De Falco E, Carnevale R. 2018. Oxidative stress in the pathogenesis of systemic scleroderma: an overview. J Cell Mol Med 22: 3308–3314.

Wang B, Guo H, Yu H, Chen Y, Xu H, Zhao G. 2015. Artesunate sensitizes ovarian cancer cells to cisplatin by downregulating RAD51. Cancer Biol Ther 16: 1548–1556.

Waseem M, Imtiaz A, Alexander A, Graham L, Contreras-Galindo R. 2025. Crosstalk between oxidative stress, mitochondrial dysfunction, chromosome instability, and the activation of the cGAS-STING/IFN pathway in systemic sclerosis. Ageing Res Rev 110: 102812.

Wei J, Bhattacharyya S, Varga J. 2011. Fibrosis in systemic sclerosis: emerging concepts and implications for targeted therapy. Autoimmun Rev 10: 267–275.

Wu X, Gao Y, Liu J, Xu X, Li Y, Wang J, Zhang H, Zhou J, Wang H. 2023. Ficolin B secreted by alveolar macrophage exosomes exacerbates bleomycin-induced lung injury via ferroptosis through the cGAS-STING pathway. Cell Death Dis 14: 577.

Yoshizaki A, Yanaba K, Iwata Y, Komura K, Ogawa A, Akiyama Y, Muroi E, Hara T, Ogawa F, Takenaka M, et al. 2010. Cell adhesion molecules regulate fibrotic process via Th1/Th2/Th17 balance in a bleomycin-induced scleroderma model. J Immunol 185: 2502–2515.

Yu S, Chen Y, Li W, Zhang X, Li H, Chen J, Chen X, Wang Y, Yang Y, Zhao J, et al. 2025. Cyclic GMP-AMP synthase expression is enhanced in systemic sclerosis-associated interstitial lung disease and stimulates inflammatory myofibroblast activation. Eur Respir J 66: 2401564.

Zeitlin SG, Baker NM, Chapados BR, Soutoglou E, Wang JYJ, Berns MW, Cleveland DW. 2009. Double-strand DNA breaks recruit the centromeric histone CENP-A. Proc Natl Acad Sci USA 106: 15762–15767.

Zhang CZ, Spektor A, Cornils H, Francis JM, Jackson EK, Liu S, Meyerson M, Pellman D. 2015. Chromothripsis from DNA damage in micronuclei. Nature 522: 179–184.

Zhang J, Wang Y, Wang H, Zhang Y, Li J, Li Y, Liu L. 2023. The role of cGAS-STING signaling in pulmonary fibrosis and its therapeutic potential. Front Immunol 14: 1273248.

